# Neural and behavioral correlates of episodic memory are associated with temporal discounting in older adults

**DOI:** 10.1101/720250

**Authors:** Karolina M. Lempert, Dawn J. Mechanic-Hamilton, Long Xie, Laura E.M. Wisse, Robin de Flores, Jieqiong Wang, Sandhitsu R. Das, Paul A. Yushkevich, David A. Wolk, Joseph W. Kable

## Abstract

When facing decisions involving trade-offs between smaller, sooner and larger, delayed rewards, people tend to discount the value of future rewards. There are substantial individual differences in this tendency toward temporal discounting, however. One neurocognitive system that may underlie these individual differences is episodic memory, given the overlap in the neural circuitry involved in imagining the future and remembering the past. Here we tested this hypothesis in older adults, including both those that were cognitively normal and those with amnestic mild cognitive impairment (MCI). We found that performance on neuropsychological measures of episodic memory retrieval was associated with temporal discounting, such that people with better memory discounted delayed rewards less. This relationship was specific to episodic memory and temporal discounting, since executive function (another cognitive ability) was unrelated to temporal discounting, and episodic memory was unrelated to risk tolerance (another decision-making preference). We also examined cortical thickness and volume in medial temporal lobe regions critical for episodic memory. Entorhinal cortical thickness was associated with reduced temporal discounting, with episodic memory performance partially mediating this association. The inclusion of MCI participants was critical to revealing these associations between episodic memory and entorhinal cortical thickness and temporal discounting. These effects were larger in the MCI group, reduced after controlling for MCI status, and statistically significant only when including MCI participants in analyses. Overall, these findings suggest that individual differences in temporal discounting are driven by episodic memory function, and that a decline in medial temporal lobe structural integrity may impact temporal discounting.

## 1. Introduction

People often have to decide between smaller/sooner and larger/later rewards. In these intertemporal choices (Strotz, 1956), individuals tend to devalue, or discount, future outcomes. This leads them to prefer immediate rewards even to larger delayed ones (Mazur, 1987), a tendency known as temporal discounting. While most individuals exhibit some degree of temporal discounting, people vary widely in the extent to which they discount delayed rewards (Peters & Büchel, 2011). Steep temporal discounting (i.e., overvaluing the present) has been linked with problematic behaviors, such as smoking (Bickel, Yi, Kowal, & Gatchalian, 2008; Yi & Landes, 2012), alcohol abuse (Vuchinich & Simpson, 1998), gambling (Reynolds, 2006), drug addiction (MacKillop et al., 2011), and excessive credit card borrowing (Meier & Sprenger, 2010). Although there is a substantial literature examining neural correlates of value at the time of intertemporal choice (Frost & McNaughton, 2017; Kable & Glimcher, 2007), the neurocognitive mechanisms underlying *individual differences* in temporal discounting remain unclear. Here we test whether individual differences in temporal discounting are linked to variability in episodic memory abilities and to variability in the structure of the medial temporal lobe (MTL), a critical neural substrate for episodic memory. To do this, we leverage variability in cognitive abilities and brain structure among older adults, including both cognitively normal older adults – that is, those who show no evidence of cognitive impairment – and those with mild cognitive impairment (MCI).

Our hypothesis that the episodic memory system may support more future-oriented intertemporal choice is motivated by a substantial body of research showing that the neural mechanisms underlying episodic memory retrieval overlap with those involved in episodic future thinking (Addis, Wong, & Schacter, 2007; Schacter & Addis, 2007). Further, people who can remember the past in more vivid detail are more likely to imagine the future more concretely (Addis, Roberts, & Schacter, 2011; Hassabis, Kumaran, Vann, & Maguire, 2007), and imagining future outcomes more vividly and concretely enhances the value of these outcomes (Rick & Loewenstein, 2008; Trope & Liberman, 2003). Supporting this idea, imagining positive future events or retrieving positive autobiographical memories prior to intertemporal choice decreases temporal discounting in young adults (Benoit, Gilbert, & Burgess, 2011; Lempert, Speer, Delgado, & Phelps, 2017; Peters & Büchel, 2010).

However, a clear association between episodic memory abilities and temporal discounting across individuals has not yet been established. There is some evidence that individuals in at-risk states for dementia, such as subjective cognitive impairment (Hu et al., 2017) and MCI (Lindbergh, Puente, Gray, Mackillop, & Miller, 2014), have higher temporal discounting rates, though these differences have not been found consistently (Chiong et al., 2016; Coelho et al., 2016). Two studies found no correlation between episodic memory and temporal discounting in older adults (Boyle et al., 2012a; Seinstra, Grzymek, & Kalenscher, 2015), but these studies (1) included only cognitively normal individuals, who might have limited variance in episodic memory abilities, and (2) used memory measures that are less susceptible to age-related decline (e.g., associative recognition; Howard, Bessette-Symons, Zhang, & Hoyer, 2006). Here we examined the relationship between episodic memory and temporal discounting in a large and well-characterized older adult sample with substantial variability in episodic memory abilities, including individuals with amnestic or multi-domain MCI, using delayed free recall measures that are sensitive to age-related decline.

In addition to relating behavioral measures of episodic memory function to temporal discounting, we also examined neuroanatomical measures of episodic memory function. Specifically, we examined the relationship between temporal discounting and cortical thickness in MTL subregions, since the MTL is critical for episodic memory (Moscovitch et al., 2005), and is also especially vulnerable to structural changes with aging (Jernigan et al., 1991; Wolk et al., 2016). In younger adults, MTL gray matter volume (Owens et al., 2017; Pehlivanova et al., 2018) significantly predicts temporal discounting rates. However, this association has not been examined in older adults.

To determine the specificity of any relationship between the episodic memory system and temporal discounting, we also examined other cognitive abilities and other kinds of decisions. Any relationship between temporal discounting and episodic memory could be driven by an association between temporal discounting and cognitive abilities in general. Indeed, fluid intelligence is associated with temporal discounting in young adults (Shamosh et al., 2008), and a composite “global cognition” score is associated with temporal discounting in older adults (Boyle et al., 2012b). A non-specific association between temporal discounting and cognition would be consistent with alternative hypotheses about the mechanism underlying individual differences in temporal discounting; for example, that frontal lobe-mediated executive functions (e.g., working memory and cognitive flexibility) are critical for making more future-oriented choices because they allow individuals to flexibly inhibit prepotent responses to choose immediate rewards (McClure & Bickel, 2014; McClure, Laibson, Loewenstein, & Cohen, 2004; Wesley & Bickel, 2014). Therefore, we also examined the relationship between temporal discounting and a composite measure of executive function, including the Trail Making Test and the Digit Span test. This allowed us to determine whether any potential relationship with temporal discounting was specific to episodic memory, rather than driven by a relationship between temporal discounting and cognitive abilities more globally.

Similarly, any relationship between cognitive abilities, such as episodic memory, and temporal discounting could be driven by an association between cognitive abilities and decision-making in general. Indeed, cognitive abilities have been associated not only with temporal discounting but also with risk tolerance (Boyle, Yu, Buchman, Laibson, & Bennett, 2011; Burks, Carpenter, Goette, & Rustichini, 2009). Our hypothesis that episodic memory contributes to more future-oriented decision-making by supporting episodic future thinking predicts that episodic memory should *not* be related to decisions that do not involve delayed outcomes. Thus, we also examined the relationship between episodic memory and risk tolerance, assessed with a risky choice task. Risky choice relies on a similar cost-benefit decision-making process as intertemporal choice, but, crucially, it does not require thinking about the future. Assessing risk preferences therefore enabled us to determine whether any potential association between cognitive abilities and temporal discounting was specific to temporal discounting, or if it extended to decision-making in general.

## 2. Materials and methods

### 2.1. Participants

100 older adults (age: 72.01±6.82 years, range: 58-93 years; male/female: 42/58, White/Black/Multi-racial: 62/36/2) completed the study. This sample was demographically representative of the greater Philadelphia area, which is 61% White, 20% Black, and 2% multi-racial (Data USA, 2020). To ensure sufficient variability in our cognitive measures, we included individuals with MCI. MCI is a syndromic label often conceptualized as an intermediate stage of cognitive decline between normal cognitive aging and mild dementia. While about 50% of MCI patients likely have underlying Alzheimer’s Disease pathology (Vos et al., 2015), the category is heterogeneous and not indicative of a specific pathological process. All subjects are part of the Clinical Core cohort of the University of Pennsylvania Alzheimer’s Disease Core Center. Given the constraint that we would only recruit from this well-characterized cohort, we selected 100 as our target sample size, in order to detect a modest correlation (*r* = 0.3) with 90% power. This study was approved by the Institutional Review Board of the University of Pennsylvania, and all participants provided informed consent. Choice task data were collected from 1/5/2017 to 2/14/2018, and all participants completed the National Alzheimer’s Coordinating Center Uniform Data Set 3.0 neuropsychological test battery (Morris et al., 2006; https://www.alz.washington.edu/WEB/data_descript.html) within one year of completing the choice tasks (87.76±70.62 days; range: 0 – 315 days). All subjects were deemed cognitively normal (*n* = 74; average Mini-Mental State Examination (MMSE) score: 29.06±1.10) or as having amnestic single- or multi-domain MCI (*n* = 26; MMSE: 27.04±2.20) based on consensus conference diagnosis by Alzheimer’s Disease clinical experts. Individuals with MCI either had a single domain of impairment, memory (*n* = 6), or were impaired in memory and at least one other domain (e.g., language or executive function; *n* = 20). As of this point (10/24/2019) based on annual evaluations, seven of the MCI participants have converted to dementia, and one of the cognitively normal participants has converted to MCI. These changes in diagnosis were recorded 335±179 days (range: 131 – 671 days) after choice task data were collected.

### 2.2. Procedure

Participants completed choice tasks assessing temporal discounting and risk tolerance (details below). The order of the tasks was counterbalanced across subjects. Both tasks were computerized (programmed in E-Prime 2.0, Psychology Software Tools, Sharpsburg, PA). Subjects were given extensive instructions as well as practice trials to confirm that they understood the tasks fully. They were also instructed that their choices were incentive-compatible. That is, at the end of the session, one choice from either the intertemporal choice or risky choice task was randomly selected to determine a bonus. Since participants did not know which choice would count, their best strategy was to treat each one as if it were the one that counted. The bonus was paid using a pre-paid debit card (Greenphire Clincard) on the day the payment was due. Because all payments were made this way, we introduced no differences in transaction costs for different types of payments (risky choice task payment, intertemporal choice immediate payment or intertemporal choice delayed payment). For delayed payments, subjects received payment on their Clincard on the date corresponding to the delay for the chosen option. The procedure lasted approximately 15 minutes. Both choice tasks were self-paced, and participants had up to 20 s to respond on each trial.

#### 2.2.1. Intertemporal choice task

On each of the 51 trials in this task, participants chose between a small amount of money available immediately, and a larger amount of money available at a specified delay (Pehlivanova et al., 2018; Senecal, Wang, Thompson, & Kable, 2012; Yu et al., 2017). The delayed outcome was always one of three amounts ($25, $30, $35). Immediate reward amounts varied from $10 − $34, and delays ranged from 1-180 days. The immediate and delayed options alternated sides of the screen randomly from trial to trial. After participants selected their choice, a checkmark appeared on the screen indicating which side they had pressed.

These 51 trials were designed as follows (Senecal et al., 2012). The goal was to capture seventeen hyperbolic discount rates, ranging from 0.0001 to 0.2525, equally distant in log-space. For each of these discount rates, three trials were drawn from its “indifference curve” – one for each of the future amounts: $25, $30, and $35. Delays were quasi-randomly generated so that the immediate amounts were integers and delays did not exceed 180 days. Like the well-validated Kirby monetary choice questionnaire (Kirby, Petry, & Bickel, 1999), this task can capture a wide range of discount rates, but with even finer gradations (Lempert, Steinglass, Pinto, Kable, & Simpson, 2018).

#### 2.2.2. Risky choice task

On each trial of this task (60 choices), participants chose between a small amount of money ($1-$68) available for certain, and a larger amount of money ($10-$100) available with some risk. All risky options entailed a 50% chance of the larger amount and a 50% chance of $0. We used a 50% probability for all trials to minimize the confounding factor of probability distortion, which might also vary among individuals. Probabilities (either 100% or 50%/50%) were displayed graphically, using pie charts. The risky and safe options alternated sides of the screen randomly from trial to trial. As with the intertemporal choice task above, the monetary amounts were selected in order to sample a range of risk tolerance parameters, with the constraint that amounts were always integers and did not exceed $100. If a participant chose the risky option on the trial randomly selected for payment, the experimenter flipped a coin to determine if they would receive the larger amount or $0.

### 2.3. Episodic memory measures

Scores on three episodic memory tasks – Word List Memory Delayed Recall, Craft Story Delayed Recall, and Benson Complex Figure Delayed Recall – were transformed to *z*-scores and averaged to form an episodic memory composite score. A composite score was used since this is more reliable than any individual measure on its own, and we had no *a priori* reason to believe that any of the individual episodic memory measures would be more strongly associated with temporal discounting than any others. All three episodic memory measures reflect recollection, rather than familiarity; the scores reflect the number of memoranda generated by participants.

#### 2.3.1. Word List Memory Delayed Recall

In this test (Morris et al., 1989), participants were presented with a list of 10 high-frequency words one at a time and were asked to read the words aloud (2 second presentation). The word list was presented 3 consecutive times, in randomized order. After every presentation, participants were asked to recall the words (Immediate Recall). After a short delay of approximately 5 minutes, the participant was asked to recall as many of the ten words as they could. We included this Delayed Recall score as one of our measures of episodic memory. Finally, participants were asked to identify the target words from a list of 10 presented words and 10 distractor words. Because performance on this recognition task was at ceiling across our sample (maximum score = 20; mean score = 19.38), we did not include it in our composite score.

#### 2.3.2. Craft Story Delayed Recall

The Craft Story 21 (Craft et al., 1996) is a paragraph story learning and recall test, similar to the Logical Memory subtest of the Wechsler Memory Scales (Wechsler, 1987). The examiner read a story aloud once, then asked the participant to repeat the details of the story in the same words read by the examiner or in their own words. Points for verbatim (exact content words) and paraphrase recall (similar contextual story units) were summed individually. After ∼15 minutes (14.52±2.30 min), the participant was asked to recall the story again. Once again, points for verbatim and paraphrase recall were summed individually. If the subject recalled no items from the Craft Story after the delay, the examiner provided a cue (“It was a story about a boy”). For this study, only the delayed paraphrase recall score (range: 1 to 25) was included in the composite score.

#### 2.3.3. Benson Complex Figure Delayed Recall

In this assessment of visuospatial memory (Possin, Laluz, Alcantar, Miller, & Kramer, 2011), participants were first asked to copy a complex figure (a simplified version of the Rey-Osterrieth complex figure), and then to draw it from memory approximately 10-15 minutes later. Their recall score was based on the accuracy and placement of reproduced elements present in the figure drawing. We used this recall score (range: 0 to 17) as our third measure of episodic memory.

### 2.4. Executive function measures

Scores on the Digit Span Backwards and Trails B-A (described below) were transformed to *z*-scores and averaged to form an executive function composite score.

#### 2.4.1. Digit Span Task (Backwards condition)

The Digit Span Task (Wechsler, 1997), part of the Wechsler Adult Intelligence Scale, is among the most widely used neuropsychological tests. It assesses auditory attention and the maintenance and manipulation of information in short-term memory. In the Forwards condition, participants are asked to repeat, in the same order, a series of digits that are read aloud to them. In the Backwards condition, participants are asked to repeat the digits in the reverse order. Because the Backwards condition is a more sensitive measure of executive function (Groeger, Field, & Hammond, 1999), the number of correct trials in this condition was used as one measure of executive function.

#### 2.4.2. Trail Making Test (Trails B-A)

The Trail Making Test (Reitan, 1992) is given in two parts, A and B. Part A involves drawing a line connecting consecutive numbers from 1 to 25 (the numbers are scattered randomly on a page) as quickly as possible. Part B involves drawing a line connecting alternating numbers and letters in sequence as quickly as possible (i.e., 1-A-2-B, etc.). The time to complete each “trail” is recorded. The difference between Trails B and Trails A times is a widely used neuropsychological measure of executive function (Davidson, Gao, Mason, Winocur, & Anderson, 2008; Stuss et al., 2001). Trails B performance involves attention, cognitive flexibility (Kortte, Horner, & Windham, 2002), and set-shifting, while Trails A performance accounts for any visuomotor or processing speed differences between subjects. Because the response time difference distribution is skewed, “Trails B-A” scores were natural log-transformed before they were *z*-scored. Trails B-A *z*-scores were also negated before being averaged in the executive function composite, since higher Trails B-A scores are indicative of worse performance.

### 2.5. Reliability and construct validity of neuropsychological measures

Given that we were comparing the effects of two composite scores that contained different measures, we wanted to ensure that these two sets of measures (1) did not significantly differ from each other in their test-retest reliability, and (2) did tap into two distinct, non-overlapping constructs.

We took advantage of the fact that participants performed these neuropsychological tests on an annual basis in order to calculate one-year test-retest reliability for each of the five measures. For this calculation, we included only cognitively normal participants, since participants with MCI are more likely to show changes in their cognitive performance from year to year. We excluded three participants because they either took the neuropsychological test battery only once (*n* = 1) or because they had only two sessions that were more than 500 days apart (*n* = 2). For the majority of participants, we took scores from the current session and compared them to those of a *previous* testing session. For participants for whom the current session was the *first* session (*n* = 6), and for those for whom the previous session was more than 500 days before the current session (*n* = 4), the scores from the *following* session were used for comparison. This left 71 participants in this analysis, with an average length of time between visits of 380.75 days (SD = 30.83 days; range [331, 483]). We computed the episodic memory composite and executive function composite for both time points, and conducted Pearson correlations between them. The correlation coefficient was 0.73 for the executive function composite, and 0.65 for the episodic memory composite. Thus, our measures had comparable and (relatively high) test-retest reliability.

Next we tested whether the five measures grouped into two distinct clusters as we proposed. We ran partial correlations between every pair of measures, controlling for every other measure. The results of these analyses can be found in Table 1. Notably, the only significant partial correlations were between the two executive function measures (*r* = 0.53) and among the three episodic memory measures (Word List Memory Delayed Recall and Benson Complex Figure Delayed Recall: *r* = 0.37; Word List Memory Delayed Recall and Craft Story Delayed Recall: *r* = 0.45; Benson Complex Figure Delayed Recall and Craft Story Delayed Recall: *r* = 0.34), with no significant relationships between executive function and episodic memory measures.

**Table 1.** Partial correlation matrix for cognitive measures.

|  | (Negative)<br>Trails B – A | Digit Span<br>Backwards | Word List<br>Memory<br>Delayed Recall | Benson<br>Complex Figure<br>Delayed Recall | Craft Story<br>Delayed Recall |
| --- | --- | --- | --- | --- | --- |
| (Negative)<br>Trails B – A | 1.00 |  |  |  |  |
| Digit Span<br>Backwards | 0.53*** | 1.00 |  |  |  |
| Word List<br>Memory<br>Delayed Recall | 0.03 | 0.05 | 1.00 |  |  |
| Benson<br>Complex Figure<br>Delayed Recall | 0.17 | -0.11 | 0.37*** | 1.00 |  |
| Craft Story<br>Delayed Recall | 0.09 | -0.02 | 0.45*** | 0.34** | 1.00 |

Note: \*\*\* $p < 0.001$ ; \*\* $p < 0.01$ . Partial correlation coefficients between each pair of measures (controlling for all four other measures) are shown.

### 2.6. Data analysis

Participants’ individual choice data for the intertemporal and risky choice tasks were fit with the following logistic function using maximum likelihood estimation:

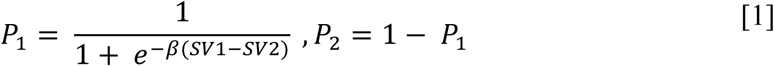

where P_1_ refers to the probability that the participant chose option 1, and P_2_ refers to the probability that the participant chose option 2. SV_1_ and SV_2_ refer to the participant’s estimated subjective value of option 1 and option 2 respectively. The scaling factor β was fitted for each individual task.

In the risky choice task, P_1_ was the probability of choosing the risky option. SV_1_ and SV_2_ (for the risky option and safe option, respectively) were estimated using a power utility function:

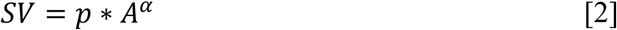

Here *A* is the amount that could be received, *p* is the probability of receipt (*p* = .5 for the risky option, *p* = 1 for the certain option), and α is a risk tolerance parameter that varies across subjects. Higher α indicates greater risk tolerance (less risk aversion).

In the intertemporal choice task, P_1_ was the probability of choosing the delayed option, and the subjective values of the options were estimated using a hyperbolic discounting function (Green & Myerson, 2004; Mazur, 1987):

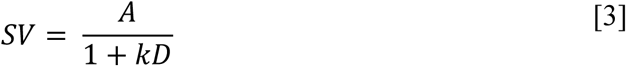

Here *A* is the amount received, *D* is the delay until receipt (for immediate rewards, *D* = 0), and *k* is a discount rate parameter that varies across subjects. Higher *k* indicates higher discounting (less tolerance of delay). Since *k* and α were not normally distributed, these values were natural log-transformed before conducting statistical analyses.

To remove any potential influence of risk tolerance on the estimates of discount rates, we also estimated discount rates using a utility-transformed hyperbolic function (Lopez-Guzman, Konova, Louie, & Glimcher, 2018):

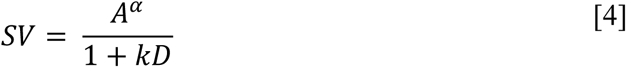

Here α is the risk tolerance parameter derived from the risky choice task. However, as we observed similar results with both sets of discount rate estimates, we only report the estimates from the linear-utility hyperbolic model (Equation [3]).

First, we examined which of our measures of interest showed age-related changes, by conducting a Pearson correlation between age and (1) temporal discounting, (2) risk tolerance, (3) episodic memory composite score, and (4) executive function composite score. To better isolate pure age effects, MCI status was included as a covariate in these analyses.

For the main analysis, we performed two multiple linear regressions, for the temporal discount rate *k* and the risk tolerance parameter α. The independent variables of interest were the episodic memory composite score and the executive function composite score. In each regression, age, sex (0 = male; 1 = female), and years of education were entered as covariates of no interest. Partial Pearson correlation coefficients and 95% confidence intervals are reported. In these regression analyses, and any analyses that included the executive function composite measure, four participants were excluded for being outliers on the Trail Making Test: one did not complete Trail Making Test Part B in the allotted time, and three had Trail Making Test Part B completion times that were more than 3 SD > mean (times of 280 s, 300 s, and 300 s).

We also ran a series of regressions to see if MCI status on its own (1 = MCI; 0 = cognitively normal; controlling for age, sex, and years of education) predicted temporal discounting, risk tolerance, and performance on episodic memory and executive function composite measures. We then examined whether our effects were driven by inclusion of MCI participants, by conducting our primary multiple linear regressions again, this time including MCI status as an additional covariate. Finally, we examined associations between episodic memory and discounting, as well as between executive function and discounting, separately in the two subgroups of participants (cognitively normal and MCI).

### 2.7. Structural MRI data acquisition and analysis

Ninety-two participants in the sample also underwent MRI scanning. Most (*n* = 53) completed the choice tasks on the same day as their structural MRI scanning session, but a subset completed the tasks at a different time. For those 39 participants, MRI scans were acquired on average 345.79±167.76 days (range: 23 – 637 days) from the choice task session.

MRI data were obtained on a Siemens Prisma 3T MRI with a 64-channel head coil. T1-weighted high-resolution magnetization-prepared rapid-acquisition gradient echo (MPRAGE; 0.8 × 0.8 × 0.8 mm^3^ voxels; TR/TE/TI=1600/3.87/950 ms; flip angle=15°) anatomical scans were collected. The medial temporal lobe subregions were segmented using an automatic pipeline, ASHS–T1 (Xie et al., 2019)^1^. This technique uses a multi-atlas label fusion approach (Wang et al., 2013) together with a tailored segmentation protocol to take into account anatomical variability in MTL cortex. It also explicitly labels dura, which has similar appearance to gray matter in T1-weighted MRI, resulting in more accurate segmentation of MTL cortex compared to other T1-MRI segmentation pipelines, such as FreeSurfer (Xie et al., 2019). In addition to a volume measure of the hippocampus, we obtained measures of mean cortical thickness in the following regions-of-interest (ROIs): entorhinal cortex, perirhinal cortex subregions Brodmann areas 35 and 36 (BA35 and BA36), and parahippocampal cortex. Cortical thickness measures used a graph-based multi-template thickness analysis pipeline taking the ASHS-T1 automatic segmentation as input (Xie et al., 2017).

In all of the regression analyses described below, measures were averaged across hemispheres. With the exception of the initial analysis of age effects, age, sex, and years of education were entered as covariates of no interest. For the hippocampal volume analyses, we included intracranial volume as an additional covariate, since volume measures are more biased by total intracranial volume than mean cortical thickness measures are (Barnes et al., 2010). For the analyses including BA35 and BA36 measures, one participant was excluded because segmentation in these regions did not pass quality control in either hemisphere. In the few cases (*n* = 7) where segmentation could be performed on one side but image quality was inadequate on the other, the mean was replaced with the thickness measure from the available side. Partial Pearson correlation coefficients and 95% confidence intervals are reported.

Our analyses of the neuroanatomical data paralleled those of the behavioral data. First, we examined which of our thickness/volume measures showed age-related changes by conducting partial correlations between thickness/volume in each of the five ROIs and age (controlling for sex and MCI status for thickness measures, and for sex, intracranial volume, and MCI status for the hippocampal volume measure). Then, to confirm that MTL structural integrity was associated specifically with episodic memory function, we examined the relationships between thickness/volume in each of the five ROIs and (1) episodic memory and (2) executive function.

For our primary question regarding the relationship between MTL structural integrity and decision-making, we conducted five regression analyses, one for each ROI, examining the relationship between its thickness/volume and temporal discounting rate. We repeated this process with the risk tolerance parameter α as the dependent variable.

If we found that structural measures of any of the MTL ROIs were significantly associated with discount rate, we planned to conduct a mediation analysis (Baron & Kenny, 1986), to see if that association was mediated by episodic memory ability. We used a Sobel test (Sobel, 1982) to test for the presence of an indirect effect (specifically, that MTL structural integrity influences discount rate via episodic memory ability), and we estimated the proportion of the total effect that could be explained through the indirect effect, along with a 95% confidence interval (Hicks & Tingley, 2011). We also planned to conduct an additional regression including *all* MTL structural measures as independent variables with temporal discounting as the dependent variable, to see whether any ROIs that significantly predicted discounting on their own remained significant predictors after controlling for thickness/volume in other MTL ROIs.

We ran a series of regressions to test if MCI status predicted thickness/volume in each of the MTL ROIs. We then planned to examine whether any neuroanatomical effects were driven by inclusion of MCI participants, by including MCI status as an additional covariate in any regressions that yielded significant effects of MTL subregion thickness/volume. Finally, we planned to examine associations between thickness/volume in MTL ROIs and temporal discounting separately in the two subgroups of participants, for any ROIs that were significant predictors of discount rate.

## 3. Results

### 3.1. Episodic memory, but not executive function, is associated with temporal discounting

100 older adults completed the National Alzheimer’s Coordinating Center Uniform Data Set 3.0 neuropsychological testing battery, as well as an intertemporal choice task and a risky choice task. See Table 2 for sample characteristics. Within this sample, when controlling for MCI status, there was no association between age and temporal discounting (*r* = 0.03; 95% CI [−0.17, 0.23]; *p* = 0.767), risk tolerance (*r* = 0.05; 95% CI [−0.15, 0.25]; *p* = 0.591), or executive function (*r* = 0.01; 95% CI [−0.19, 0.21]; *p* = 0.923), although increased age was associated with worse episodic memory (*r* = −0.25; 95% CI [−0.43, −0.06]; *p* = 0.014). In all analyses reported below, age, sex, and years of education were entered as covariates, and partial correlation coefficients are reported.

**Table 2.** Characteristics of participants (*N* = 100)

| Characteristic | CN ($n = 74$)<br>Mean (SD, Range) or % | MCI ($n = 26$)<br>Mean (SD, Range) or % |
| --- | --- | --- |
| Age | 72.07 (6.83, 58-93) | 71.85 (6.93, 58-87) |
| Sex | 66% Female, 34% Male | 35% Female, 65% Male |
| Race | 58% White, 39% Black,<br>3% Multi-racial | 73% White, 27% Black |
| Years of education | 15.49 (2.80, 9-20) | 17.31 (2.46, 12-20) |
| Cognitive measures: Raw scores | Mean (SD, Range) | Mean (SD, Range) |
| Word List Memory Delayed Recall<br>(# words recalled) | 8.61 (1.57, 3-10) | 4.35 (2.42, 0-10) |
| Craft Story Delayed Recall (# story<br>units recalled) | 16.04 (3.56, 6-23) | 6.77 (5.89, 0-17) |
| Benson Complex Figure Delayed<br>Recall | 11.72 (2.66, 4-16) | 4.85 (4.55, 0-14) |
| Digit Span Backwards (# correct) | 6.95 (2.21, 2-13) | 6.12 (1.61, 3-9) |
| Trail Making Test (Part B minus<br>Part A) RT | 42.59 s (21.76 s, 10-126 s) | 60.14 s (32.29 s, 17-153 s)* |

Note: CN = cognitively normal; MCI = mild cognitive impairment; RT = reaction time. \* $n = 4$ participants excluded for not completing Trail Making Test Part B in the allotted time ( $n = 1$ ) or for having a reaction time on Trail Making Test Part B that was more than 3 SD > mean ( $n = 3$ ; 280 s, 300 s, and 300 s).

Episodic memory, but not executive function, was associated with temporal discounting rate. When entering both the episodic memory composite score and executive function composite score as independent variables in a regression predicting temporal discounting rate, better performance on episodic memory tasks was associated with reduced temporal discounting (*r* = −0.32; 95% CI [−0.49, −0.12]; *p* = 0.002; *n* = 96; Fig. 1A), while the association between executive function and temporal discounting was not significant (*r* = 0.08; 95% CI [−0.13, 0.28]; *p* = 0.461; Fig. 1B). That is, individuals with better episodic memory tended to discount delayed rewards less, even after controlling for variance that could be explained by executive function. When conducting separate regressions for episodic memory and executive function (still controlling for age, sex, and years of education), there was similarly a significant association between episodic memory and temporal discounting (*r* = −0.30; 95% CI [−0.47, −0.11]; *p* = 0.003; *n* = 100), but not between executive function and temporal discounting (*r* = −0.04; 95% CI [−0.24, 0.16], *p* = 0.693; *n* = 96).

**Fig. 1.**
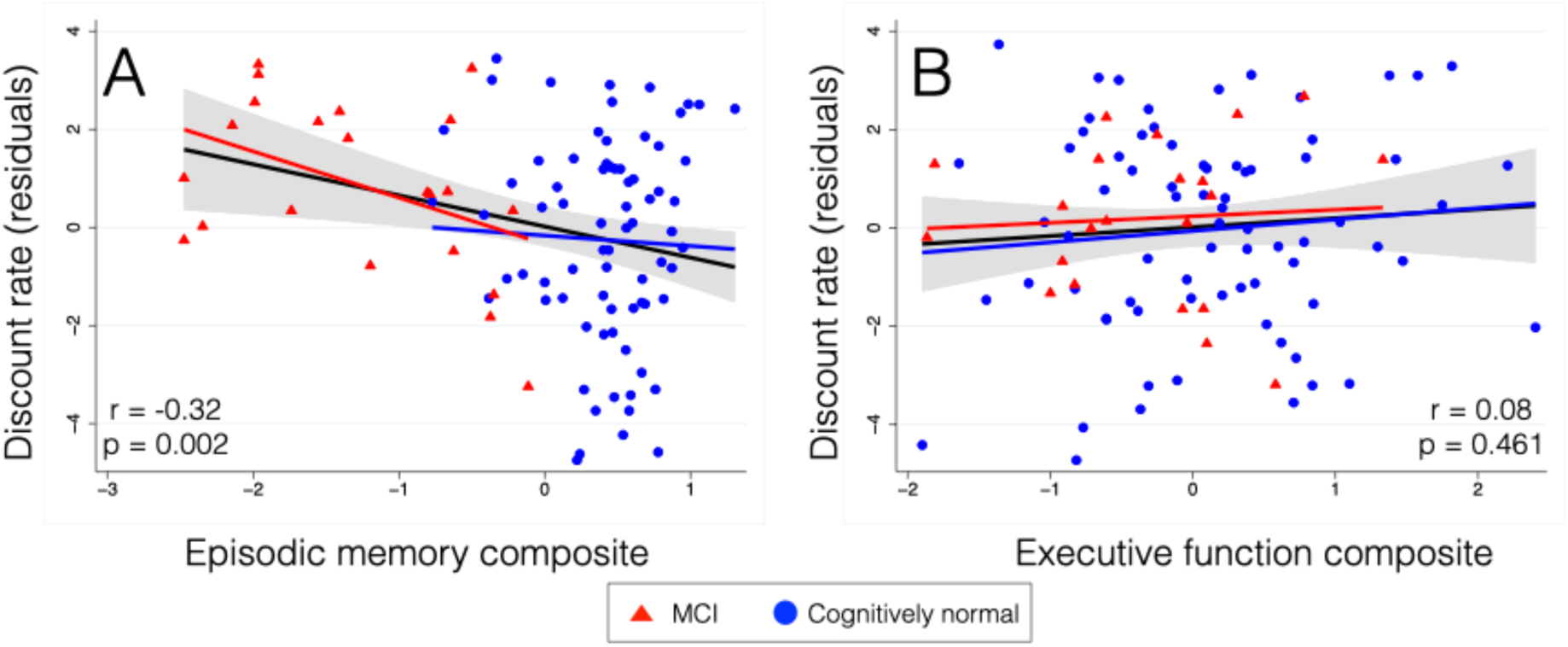
Associations between cognitive measures and temporal discounting. The episodic memory composite measure (A) is significantly correlated with temporal discounting rate: people with better memory are more patient for future rewards. The residual discount rate after adjusting for age, sex, years of education, and executive function is shown. Performance on the executive function composite measure (B) is not significantly correlated with discount rate. The residual discount rate after adjusting for age, sex, years of education, and episodic memory is shown. MCI = Mild Cognitive Impairment. The linear fit for data from the whole sample along with a 95% confidence interval is overlaid on the plots in black. Linear fits for the cognitively normal participants are overlaid in blue and for MCI participants in red.

This association between episodic memory measures and temporal discounting was specific to discounting, and did not extend to decision tendencies in the risk domain. From our risky choice task, we derived a measure of an individual’s risk tolerance by assuming a power function for utility and estimating a risk tolerance parameter α. There was no relationship between the episodic memory composite score and risk tolerance or between the executive function composite score and risk tolerance, whether both composite scores were entered as predictors in the regression together (episodic memory *r* = 0.07; 95% CI [−0.14, 0.27]; *p* = 0.486; executive function *r* = 0.08; 95% CI [−0.13, 0.28]; *p* = 0.466) or separately (episodic memory *r* = 0.10; 95% CI [−0.10, 0.29]; *p* = 0.350; executive function *r* = 0.11; 95% CI [−0.10, 0.31]; *p* = 0.297).

We recruited individuals with MCI into our sample in order to increase the range of episodic memory scores. As expected, when controlling for age, sex, and years of education, MCI status (0 = cognitively normal; 1 = MCI) was associated with worse performance on our episodic memory composite measure (*r* = −0.79; 95% CI [−0.85, −0.70]; *p* < 0.001). MCI participants also performed significantly worse on executive function measures (*r* = −0.27; 95% CI [−0.45, −0.07]; *p* = 0.008), suggesting that their impairment extended beyond the memory domain. There was a significant effect of MCI status on discount rate (*r* = 0.26; 95% CI [0.06, 0.43]; *p* = 0.011), with MCI participants displaying increased temporal discounting overall. Consistent with the null relationship between our cognitive measures and risk tolerance, there were no differences between MCI and cognitively normal participants in risk tolerance (*r* = − 0.11; 95% CI [−0.30, 0.09]; *p* = 0.264).

The association between episodic memory and temporal discounting was contingent on our inclusion of MCI participants. When including MCI status as an additional covariate (along with executive function, age, sex, and years of education) in the regression, MCI status was no longer a significant predictor of temporal discounting (*r* = 0.03; 95% CI [−0.18, 0.23]; *p* = 0.792), and the relationship between episodic memory and temporal discounting fell to a trend level (*r* = −0.19; 95% CI [−0.38, 0.02]; *p* = 0.078). Results were similar when the executive function predictor was excluded (episodic memory *r* = −0.17; 95% CI [−0.36, 0.03]; *p* = 0.101; MCI *r* = 0.03; 95% CI [−0.17, 0.23]; *p* = 0.758). When running separate regressions within each diagnosis subgroup, there was no significant association between episodic memory and discount rate (controlling for executive function, age, sex, and years of education: cognitively normal, *n* = 74: *r* = −0.07; 95% CI [−0.30, 0.17]; *p* = 0.553; MCI, *n* = 22: *r* = −0.45; 95% CI [−0.76, 0.02]; *p* = 0.064), but the correlation coefficients suggest that the relationship was much stronger in the MCI group, in which there was trend-level significance. Overall, these results suggest that the association between episodic memory and temporal discounting is driven primarily by a linear relationship within the MCI sample.

### 3.2. Entorhinal cortical thickness is associated with temporal discounting

A large subset of participants (*n* = 92) had structural MRI data. We examined associations with structural measures from five medial temporal lobe subregions: cortical thickness in entorhinal cortex, BA35, BA36, and parahippocampal cortex, and hippocampal volume. Age was not associated with mean cortical thickness in entorhinal cortex (*r* = −0.20; 95% CI [−0.39, 0.01]; *p* = 0.062), BA35 (*r* = −0.19; 95% CI [−0.38, 0.02]; *p* = 0.073), or BA36 (*r* = −0.13; 95% CI [−0.33, 0.08]; *p* = 0.234), but was associated with thickness in parahippocampal cortex (*r* = −0.24; 95% CI [−0.43, −0.03]; *p* = 0.022), and hippocampal volume (*r* = −0.30; 95% CI [−0.48, −0.10]; *p* = 0.004). In all analyses reported below, age is included as a covariate of no interest, along with sex and years of education (for hippocampal volume analyses, intracranial volume is included as an additional covariate).

We confirmed that structural integrity of the medial temporal lobe was associated with episodic memory, but not executive function, in our dataset. Structural measures from all of these medial temporal lobe regions were associated with episodic memory (entorhinal cortex *r* = 0.54; 95% CI [0.37, 0.67]; *p* < 0.001; BA35 *r* = 0.41; 95% CI [0.22, 0.57]; *p* < 0.001; BA36 *r* = 0.28; 95% CI [0.07, 0.46]; *p* = 0.009; parahippocampal cortex *r* = 0.24; 95% CI [0.03, 0.43]; *p* = 0.024; hippocampus *r* = 0.57; 95% CI [0.41, 0.70]; *p* < 0.001). In contrast, none of the medial temporal lobe structural measures were associated with executive function (entorhinal cortex *r* = 0.07; 95% CI [−0.14, 0.27]; *p* = 0.543; BA35 *r* = 0.10; 95% CI [−0.11, 0.30]; *p* = 0.385; BA36 *r* = 0.08; 95% CI [−0.13, 0.28]; *p* = 0.466; parahippocampal cortex *r* = 0.06; 95% CI [−0.15, 0.26]; *p* = 0.586; hippocampus *r* = 0.10; 95% CI [−0.11, 0.30]; *p* = 0.352).

Given the association between behavioral measures of episodic memory and temporal discounting, next we conducted an exploratory analysis relating structural integrity in the medial temporal lobe to temporal discounting. When examining each of the five MTL subregions in separate regressions, temporal discounting was associated with mean cortical thickness in the entorhinal cortex (*r* = −0.28; 95% CI [−0.46, −0.08]; *p* = 0.008; *n* = 92; Fig. 2), but was not associated with cortical thickness in BA35 (*r* = −0.09; 95% CI [−0.29, 0.12]; *p* = 0.382; *n* = 91), BA36 (*r* = −0.18; 95% CI [−0.38, 0.03]; *p* = 0.094; *n* = 91), or parahippocampal cortex (*r* = −0.17; 95% CI [−0.36, 0.04]; *p* = 0.116; *n* = 92), or with the volume of the hippocampus (*r* = −0.06; 95% CI [−0.27, 0.15]; *p* = 0.587; *n* = 92). Cortical thickness in entorhinal cortex was also related specifically to discounting, and not to risk tolerance (*r* = 0.05; 95% CI [−0.16, 0.26]; *p* = 0.642; see Table 3 for full list of partial correlation coefficients). Finally, when examining all five MTL subregions in the same regression, entorhinal cortical thickness was associated with temporal discounting above and beyond the other medial temporal lobe thickness/volume measures (entorhinal cortex *r* = −0.26; 95% CI [−0.45, −0.05]; *p* = 0.018; parahippocampal cortex *r* = −0.13; 95% CI [−0.34, 0.09]; *p* = 0.248; BA35 *r* = 0.12; 95% CI [−0.10, 0.33]; *p* = 0.271; BA36 *r* = − 0.14; 95% CI [−0.35, 0.08]; *p* = 0.209; hippocampus *r* = 0.10; 95% CI [−0.12, 0.31]; *p* = 0.359).

**Table 3.** Associations between cortical thickness in regions of interest and choice measures.

| Region | Discount rate | Risk tolerance |
| --- | --- | --- |
| Entorhinal cortex | **-0.28\*\*[-0.46, -0.08]** | 0.05 [-0.16, 0.26] |
| BA36 | -0.18 [-0.38, 0.03] | -0.13 [-0.33, 0.08] |
| BA35 | -0.09 [-0.29, 0.12] | -0.02 [-0.23, 0.19] |
| Parahippocampal cortex | -0.17 [-0.36, 0.04] | 0.03 [-0.18, 0.24] |
| Hippocampus (volume) | -0.06 [-0.27, 0.15] | 0.12 [-0.09, 0.32] |

Note: Partial correlation coefficients, controlling for age, sex, years of education, and intracranial volume (for hippocampus only) are shown, along with 95% confidence intervals. \*\* $p < 0.01$ (uncorrected for multiple comparisons).

**Fig. 2.**
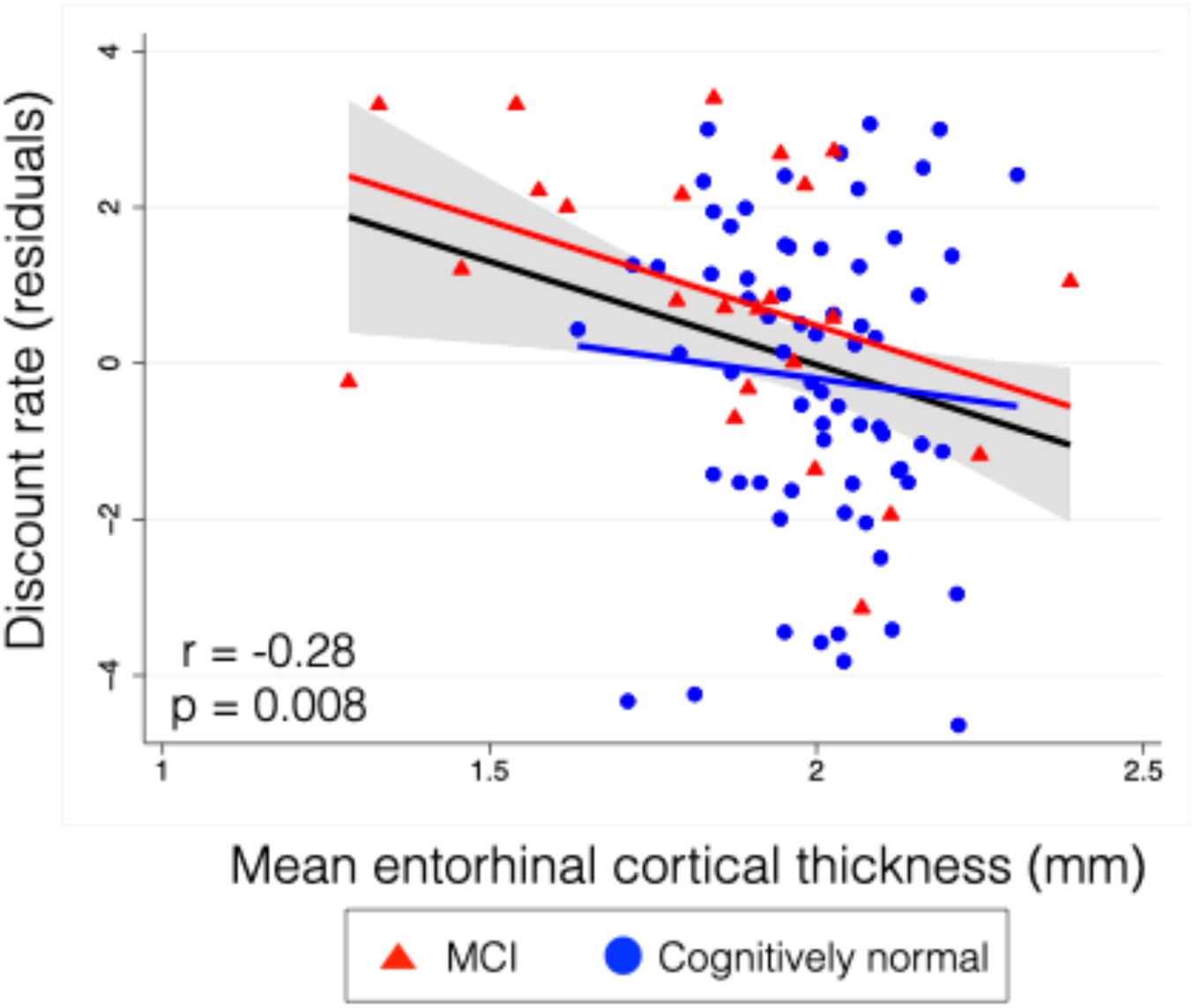
Association between entorhinal cortical thickness (in millimeters) and temporal discounting rate (*n* = 92). The entorhinal cortex was the only one of our medial temporal lobe ROIs that was significantly and robustly associated with temporal discounting rate: people with more entorhinal cortical thickness were more patient for future rewards. The residual plot after adjusting for age, sex, and years of education is shown. MCI = Mild Cognitive Impairment. The linear fit for data from the whole sample along with a 95% confidence interval is overlaid on the plot in black. The linear fit is in blue for the cognitively normal participants, and in red for the MCI participants.

Episodic memory ability partially mediated the relationship between entorhinal cortical thickness and temporal discounting. When entorhinal cortical thickness and episodic memory were entered in the same regression to predict discount rate (with age, sex, and years of education as covariates), we found that entorhinal cortical thickness no longer predicted discounting (*r* = −0.12; 95% CI [−0.33, 0.09]; *p* = 0.249). A Sobel test revealed that there was a significant indirect effect (*t* = 2.07; *p* = 0.039), whereby entorhinal cortical thickness influences temporal discounting through its influence on episodic memory. Approximately 49% (bootstrapped 95% CI [28.45%, 179.11%]) of the total effect of entorhinal cortical thickness on temporal discounting could be explained through this indirect effect.

Individuals with mild cognitive impairment showed a stronger relationship between entorhinal cortical thickness and temporal discounting. MCI status was a significant predictor of structural integrity in entorhinal cortex (*r* = −0.33; 95% CI [−0.50, −0.13]; *p* = 0.002), BA35 (*r* = − 0.26; 95% CI [−0.45, −0.05]; *p* = 0.016), BA36 (*r* = −0.22; 95% CI [−0.41, −0.009]; *p* = 0.041), and hippocampus (r = −0.41; 95% CI [−0.57, −0.22]; *p* < 0.001) but not parahippocampal cortex (*r* = − 0.17; 95% CI [−0.37, 0.04]; *p* = 0.110). When including MCI status as an additional covariate when examining the association between entorhinal cortical thickness and temporal discounting, MCI status was no longer a significant predictor of discounting (*r* = 0.19; 95% CI [−0.02, 0.38]; *p* = 0.083), and the relationship between entorhinal cortical thickness and temporal discounting was reduced, although still significant (*r* = −0.21; 95% CI [−0.40, −0.001]; *p* = 0.044). When examining the cognitively normal and MCI subgroups separately, the association was stronger in the MCI group (MCI, *n* = 24: *r* = −0.44; 95% CI [−0.73, −0.01]; *p* = 0.047; cognitively normal, *n* = 68: *r* = −0.08; 95% CI [−0.32, 0.17]; *p* = 0.506). Overall, this pattern resembles what was observed in the behavioral data, and suggests that the association between entorhinal cortical thickness and temporal discounting is strongest within the MCI sample, perhaps reflecting increased variance in MTL atrophy in this subgroup.

## 4. Discussion

In a diverse group of cognitively normal and MCI older adults, we found that better episodic memory ability was associated with reduced temporal discounting, or a relatively greater preference for larger, delayed rewards. This association was specific to episodic memory and temporal discounting, as executive function (another cognitive ability) was unrelated to temporal discounting, and episodic memory was unrelated to risk tolerance (another decision-making preference). In an exploratory analysis relating cortical thickness in the MTL to temporal discounting, we found that thickness in the entorhinal cortex was associated with temporal discounting, and that the relationship between entorhinal cortical thickness and discount rate was partially mediated by episodic memory ability. The inclusion of MCI participants was critical to revealing these behavioral and neural associations, which tended to be much stronger in this subgroup than in cognitively normal participants. In fact, the associations between episodic memory and temporal discounting, and between entorhinal cortical thickness and temporal discounting, were not statistically significant when examining the cognitively normal subgroup on its own.

To our knowledge, this is the first study to link episodic memory abilities with temporal discounting. Previous studies of intertemporal choice in individuals with MCI and Alzheimer’s Disease have yielded inconsistent results (Chiong et al., 2016; Coelho et al., 2016; Lebreton et al., 2013; Lindbergh et al., 2014). In cognitively normal older adults, one previous study found that global cognition was associated with temporal discounting, but episodic and semantic memory composite scores were not (Boyle et al., 2012a), and another also found no relationship between episodic memory ability and temporal discounting (Seinstra et al., 2015). Consistent with these two studies, we also found no relationship between episodic memory and temporal discounting within the cognitively normal subgroup in our study. However, when leveraging a large and well-characterized sample with substantial variability in episodic memory retrieval ability, including individuals with MCI, we detected a significant association between memory function and temporal discounting.

Another contribution of the current study is linking neural correlates of memory with temporal discounting. Structural integrity in the MTL has previously been associated with temporal discounting in adolescents (Pehlivanova et al., 2018) and young and middle-aged adults (Owens et al., 2017). Here we found in a heterogenous group of older adults, including those with MCI, that entorhinal cortical thickness was associated with temporal discounting. Perhaps surprisingly, hippocampal volume was not associated with temporal discounting, despite the importance of this region for the formation of context-rich episodic memories. However, entorhinal cortex was the primary region where structural differences were associated with temporal discounting in the largest previous study (Owens et al., 2017). Entorhinal cortex is also one of the earliest sites for development of the neurofibrillary tangle pathology associated with aging and Alzheimer’s Disease (Braak & Braak, 1991). The exact mechanism by which the entorhinal cortex supports future-oriented choice warrants further study. One possibility is that given its role as a “relay station” between hippocampus and prefrontal cortex (Agster & Burwell, 2009; Apergis-Schoute, Pinto, & Paré, 2006), the entorhinal cortex facilitates the modulation of valuation by episodic future thinking at the time of choice. Another possibility is that its involvement in memory for timing or duration (Lositsky et al., 2016; Montchal, Reagh, & Yassa, 2019) influences prospective duration estimates. While our analysis of MTL subregions was exploratory and preliminary, our findings are strengthened by our use of state-of-the-art methods for segmentation (which included rigorous quality checks) in a well-characterized group of older adults with variability in the structural integrity of these regions.

The inclusion of MCI participants was critical here, as there was no significant correlation between episodic memory measures and temporal discounting, or between entorhinal cortical thickness and temporal discounting, in the participants who were classified as cognitively normal. This might reflect a limited range of variability in episodic memory abilities or entorhinal cortical thickness in the cognitively normal group. It might also reflect a non-linear relationship between episodic memory or entorhinal cortical thickness and temporal discounting, such that the association is stronger after some threshold of memory decline or structural degeneration. In either case, this suggests that effects on temporal discounting may be most clinically relevant only in individuals who show memory and neuroanatomical changes predictive of incipient Alzheimer’s Disease.

We cannot definitively rule out the possibility that MCI participants have higher discount rates for other reasons related to their diagnosis (e.g., awareness of their diagnosis), and that discounting is not truly directly related to episodic memory or entorhinal cortical thickness. Several results argue against that possibility, however. First, MCI participants were also impaired on executive function tasks (albeit to a lesser degree), but temporal discounting was specifically correlated with episodic memory. Similarly, in the neuroanatomical data, MCI participants had reduced thickness/volume in most of our regions-of-interest, but temporal discounting was specifically correlated with entorhinal cortical thickness. Second, within the MCI group, the partial correlation between temporal discounting and episodic memory was high (*r* = −0.45), as was the partial correlation with entorhinal cortical thickness (*r* = −0.44); these continuous associations would be unlikely if it was participants’ awareness of their diagnosis that changed preferences. Third, there was no relationship between MCI status and discounting when controlling for episodic memory ability or entorhinal cortical thickness. Finally, the relationship between entorhinal cortical thickness and discount rate was significant even when controlling for diagnosis, and the relationship between episodic memory and discount rate remained at a trend level. Future studies using episodic memory measures that show sufficient variance in young and cognitively normal older adults will shed additional light on the generalizability of this relationship. Nevertheless, the finding that our cohort of MCI participants, a group that is defined by their memory deficits, show increased temporal discounting is evidence for the hypothesis that episodic memory drives individual differences in intertemporal preferences.

Performance on standard measures of executive function (Digit Span Backwards and Trails B – A) was not associated with temporal discounting. Thus, our findings provide key evidence that episodic memory processes are a more important contributor to future-directed decision-making than executive function. Although there is a well-documented association between temporal discounting and fluid intelligence (Burks et al., 2009; Kable et al., 2017; Shamosh et al., 2008), in principle, episodic memory could underpin this association. Recent research has shown that the strong correlation between working memory and general intelligence (Ackerman, Beier, & Boyle, 2005) may be driven by individual differences in episodic memory processes, such as search and retrieval (Healey, Crutchley, & Kahana, 2014; Mogle, Lovett, Stawski, & Sliwinski, 2008; Unsworth, Brewer, & Spillers, 2009). Furthermore, the most successful manipulations of temporal discounting to date involve activating episodic memory circuitry by encouraging people to imagine future events (Bulley, Henry, & Suddendorf, 2016). On the other hand, taxing frontal executive processes does not increase impulsive choice, but rather, decreases choice consistency (i.e., the ability to maintain consistent preferences across trials; Franco-Watkins et al., 2010; Olschewski et al., 2018). However, a couple of caveats warrant mention. First, executive function is a heterogeneous construct, and the neuropsychological measures that we used capture only some aspects of executive function (e.g., manipulation in working memory in the Digit Span Test and set-shifting in Trails B – A). Therefore, it is possible that some other aspects of executive function not measured here (e.g., flexible reasoning, inhibition, or abstraction) would show a relationship with temporal discounting. Nevertheless, our results rule out the possibility that *all* cognitive abilities are associated with temporal discounting. A second limitation is that, as our investigation focused on episodic memory, we prioritized recruitment of MCI individuals with impairments in the memory domain. Including more individuals with non-amnestic MCI could have revealed an association between temporal discounting and executive function.

We found no association between either executive function or episodic memory and risk tolerance. Previous research has linked cognitive abilities with risk preferences, including in older adults (Boyle et al., 2011). Specifically, people who are more educated (Donkers et al., 1999), have higher intelligence (Burks et al., 2009), and have better global cognition (Boyle et al., 2011; Frederick, 2005) are more risk-seeking. It is possible that other cognitive abilities not assessed here are preferentially involved in decisions under risk. It is also possible that the association between cognitive abilities and risk tolerance holds primarily when risky choices have higher expected value, which was not always the case in our choice set.

The results of the current study elucidate the inconsistent findings regarding aging and intertemporal decision-making. We did not find a significant relationship between age and temporal discounting rate, whether looking at our sample overall (*r* = 0.06, *p* = 0.56), or just within the cognitively normal (*r* = 0.03, *p* = 0.79) or MCI (*r* = 0.17, *p* = 0.41) groups. This is consistent with previous studies showing that sound economic decision-making remains intact in older adults (Li, Baldassi, Johnson, & Weber, 2013; Li et al., 2015), and that changes in economic decision-making with aging are more likely to be underpinned by changes in cognitive abilities (Henninger, Madden, & Huettel, 2010). Our results suggest specifically that temporal discounting may increase with age to the extent that episodic memory declines. However, we cannot draw this conclusion from our cross-sectional investigation. Longitudinal work has shown that changes in cognitive function in older adults are associated with concomitant changes in temporal discounting (James, Boyle, Yu, Han, & Bennett, 2015). Future research, perhaps with the same cohort used here, will reveal whether episodic memory decline has a causal influence on intertemporal decision-making.

In sum, the current study sheds light on the cognitive and neural mechanisms underlying individual differences in temporal discounting. It also contributes to our understanding of decision-making in the context of aging. These findings may aid in the development of interventions to promote greater patience in economic decisions, especially as cognition declines.

## Acknowledgements

We would like to thank Laura Saad, Arun Pilania, Joseph Harrison, and Jacqueline Lane for assistance with data collection.

## Footnotes

1 ASHS-T1 is a follow-up pipeline to ASHS (Automatic Segmentation of Hippocampal Subfields) which is used to label hippocampal subfields and MTL cortical subregions in T2-MRI. It was recently extended to T1-MRI and modified to label MTL cortical subregions and partition hippocampus into anterior and posterior regions, since hippocampal subfields are not generally distinguishable in T1-MRI.

